# Perfecting and extending the near-infrared biological window

**DOI:** 10.1101/2021.04.19.440389

**Authors:** Zhe Feng, Tao Tang, Tianxiang Wu, Xiaoming Yu, Yuhuang Zhang, Meng Wang, Junyan Zheng, Yanyun Ying, Siyi Chen, Jing Zhou, Xiaoxiao Fan, Shengliang Li, Mingxi Zhang, Jun Qian

## Abstract

*In vivo* fluorescence imaging in the second near-infrared window (NIR-II) has been considered as a promising technique for visualizing the mammals. However, the definition of the NIR-II region and the mechanism accounting for the excellent performance still need to be perfected. Herein, we simulated bioimaging in the NIR spectral range (to 2340 nm), confirmed the positive contribution of moderate light absorption by water in intravital imaging and perfected the NIR-II window as 900-1880 nm, where the 1400-1500 nm was defined as NIR-IIx region and the 1700-1880 nm was defined as NIR-IIc region, respectively. Moreover, the 2080-2340 nm was newly proposed as the third near-infrared (NIR-III) window, which was believed to provide the best imaging quality. The wide-field fluorescence microscopy in brain, in addition, was performed around NIR-IIx region with excellent optical sectioning strength and the largest imaging depth of in vivo NIR-II fluorescence microscopy to date. We also proposed 1400 nm long-pass detection in off-peak NIR-II imaging whose profits exceeded those of NIR-IIb imaging, using bright fluorophores with short peak emission wavelength.

## Introduction

Fluorescence imaging has been widely utilized in medical practices. With the deepening of understanding of the interaction between light and bio-tissue as well as the cost decline of detection technique, the fluorescence imaging wavelength as a whole is red shifted constantly from visible range to near-infrared region.^1,2^ The energy loss when light propagates in the biological media could be blamed on the absorption attenuation and the scattering disturbance. The absorption loss determines whether we could catch the signals while the scattering signals always reduce definition of images. Additionally, excessive light absorption in bio-tissue might induce tissue injury. The autofluorescence from some biomolecules is always mingled with the useful signals and eventually becomes the background of taken images. Thus, the deep-rooted beliefs that light absorption and scattering are totally harmful for fluorescence catching, urging most researchers chase a perfect window with minimal photon absorption and scattering for bioimaging. Due to the generally accepted less photon scattering, the fluorescence bioimaging in the second near-infrared window (NIR-II) gives admirable image quality, especially when deciphering the deep-buried signals *in vivo*.^3–9^ Nowadays, NIR-II fluorescence imaging has already guided complicated liver-tumor surgery in clinic.^10^ However, the constructive role of light absorption, to some extent, seems to be ignored. The final presentation of high-quality images even makes the overstated positive effect of scattering suppression by lengthening wavelength more convincing, since the absorption simultaneously is considered to attenuate the signals. As a matter of fact, some works have revealed absorption-induced resolution enhancement in the scattering media due to the depressing of long-optical-path background signals.^11,12^ Yet how to take full advantage of light absorption to select suitable fluorescence imaging window remains unspecified.

The definition of NIR-II window has been always limited to 1000-1700 nm, prompting the launch of various NIR emitters with the peak emission wavelength beyond 1000 nm^13–15^ and even beyond 1500 nm (NIR-IIb region, 1500-1700 nm) ^16–18^. Some existing and developing fluorophores with peak emission below 1000/1500 nm but bright emission tail beyond 1000/1500 nm, meanwhile, are also well suited for NIR-II/NIR-IIb fluorescence imaging.^19–24^ It must be admitted that the design and synthesis of bright and long-wavelength NIR emitters are still full of challenges nevertheless. Besides, the positive role of light absorption mentioned above in NIR-II fluorescence imaging could undermine the privilege of ultralong emitters; in other words, it is not an inevitability that the longer the imaging wavelength, the better the imaging performance. Therefore, preparation of wavelength-tunable NIR-II fluorophores with stable brightness is still not easily available, but of great significance for us to search for the optimum imaging window, even exceeding NIR-IIb region (e.g. beyond 1700 nm). In addition, organic agents are always considered well biocompatible but it is indeed hard to equip the organic dyes with both long wavelength and strong emission. Since the emission tailing usually occupies a few of the whole, the agents were directly applied to detection in some given long-pass (LP) spectral regions and perform high-contrast imaging.^25–27^ However, the past experience guides us to avoid the light absorption peak of biomolecules (such as water) when determining the imaging window. The best LP band for imaging, therefore, still needs to be carefully verified, taking the positive influence of absorption into consideration.

Microscopic examination of the tiny bio-structure is always conductive to the understanding of biological processes and diagnosis, as well as treatment for certain diseases. Fluorescence wide-field microscopy in NIR-II region has shown its excellent strength of deep deciphering in bio-tissues of rodents^9,28–30^ and even non-human primates^22,31^. The user-friendly imaging mode with high temporal resolution could assist operators monitor the dynamic process in real time such as blood flow. However, defocusing signals and scattering light are often collected along with the targeted information, and thus become the strong background, consequently reducing the image contrast. Some advanced NIR-II microscopic imaging techniques, such as confocal^18,28,29,32^ and light-sheet microscopy^33^, aim to counteract the image background via adjusting the collection and excitation pattern. However, the pinhole-introduced scanning confocal microscopy wastes the useful signals inevitably and lengthens the imaging duration compared with area-detection microscopy. The light-sheet excitation always places great demands on the transparency of sample. In this way, the microscopy with high spatio-temporal resolution, deep penetration and easy operation is still urgently needed.

Herein, we used Monte Carlo method to simulate the NIR imaging and innovatively proposed the well-performance imaging in 1400-1500 nm, 1700-1880 nm and 2080-2340 nm, which were defined as NIR-IIx, NIR-IIc and the third near-infrared (NIR-III) window, respectively. Lead sulfide (PbS) quantum dots (QDs) have exhibited the advantages of high fluorescence brightness and tunable emission wavelength^34–36^ for noninvasive high-performance in vivo imaging in the NIR-II window.^14,18,33,37,38^ We designed and synthesized a series of PbS/CdS core–shell quantum dots (CSQDs), and then hydrated them with polyethylene glycol (PEG). Assisted by the bright QDs with peak emission wavelength at ~1100 nm, ~1300 nm and ~1450 nm, we found the detection regions around the absorption peaks of water always provide vastly improved image quality and thus the definition of NIR-II window was further perfected as 900-1880 nm. The 1400-1500 nm which was defined as the near-infrared IIx (NIR-IIx) region was proved to provide more superior fluorescence image than NIR-IIb region. The *in vivo* NIR-IIx fluorescence microscopic cerebral vasculature imaging with convenient optical setup but strongly-suppressed background was conducted with admirable image contrast and the imaging depth reached to ~1.3 mm, which represented the deepest *in vivo* NIR-II fluorescence imaging in mice brain to date. Considering the emission of substantial fluorophores peaking shorter than 1400 nm but holding bright emission tailing, we proposed an effective usage of 1400 nm LP collection with better performance than NIR-IIb region. We believed these results are pretty crucial for the further development of NIR fluorescence imaging.

## Results

### Simulation and discussion on the near-infrared fluorescence imaging

General cognition of NIR-II window guides us to emphasize the scattering depression with the increase of wavelength but underestimate the constructive effect of absorption. As a matter of fact, the light absorbers would preferentially deplete the multiply scattered photons in propagation in theory since scattered photons have longer path lengths through the biological medium than ballistic photons (Scheme 1). Monte Carlo method was utilized to simulate the propagation of photons in biological tissues here^39^ and the imaged line sample was shown in Fig. S1. The scattering depression would benefit the transmission of ballistic photons and the effective collection. With the decrease of the reduced scattering coefficient (μ_s’ =_ 10 mm^−1^ in Fig. 1a; μ_s’ =_ 3 mm^−1^ in Fig. 1b; μ_s’ =_ 0.2 mm^−1^ in Fig. 1c) but the same absorption coefficient (μ_a_) of 0.3 mm^−1^, the image became clearer (Fig. 1a-c), and the full width at half height (FWHM) declined (Fig. 1d) and the signal-to-background ratio (SBR) grew significantly (Fig. 1e). Then the μ_s’_ was set to 1 mm^−1^, and μ_a_ was 0.01, 0.1, and 1 mm^−1^ in Fig. 1f, g and h, respectively. It could be seen that the light absorption in tissue could inhibit the strong background and enhance the spatial resolution (Fig. 1i) as well as imaging contrast (Fig. 1j).

**Scheme 1.**
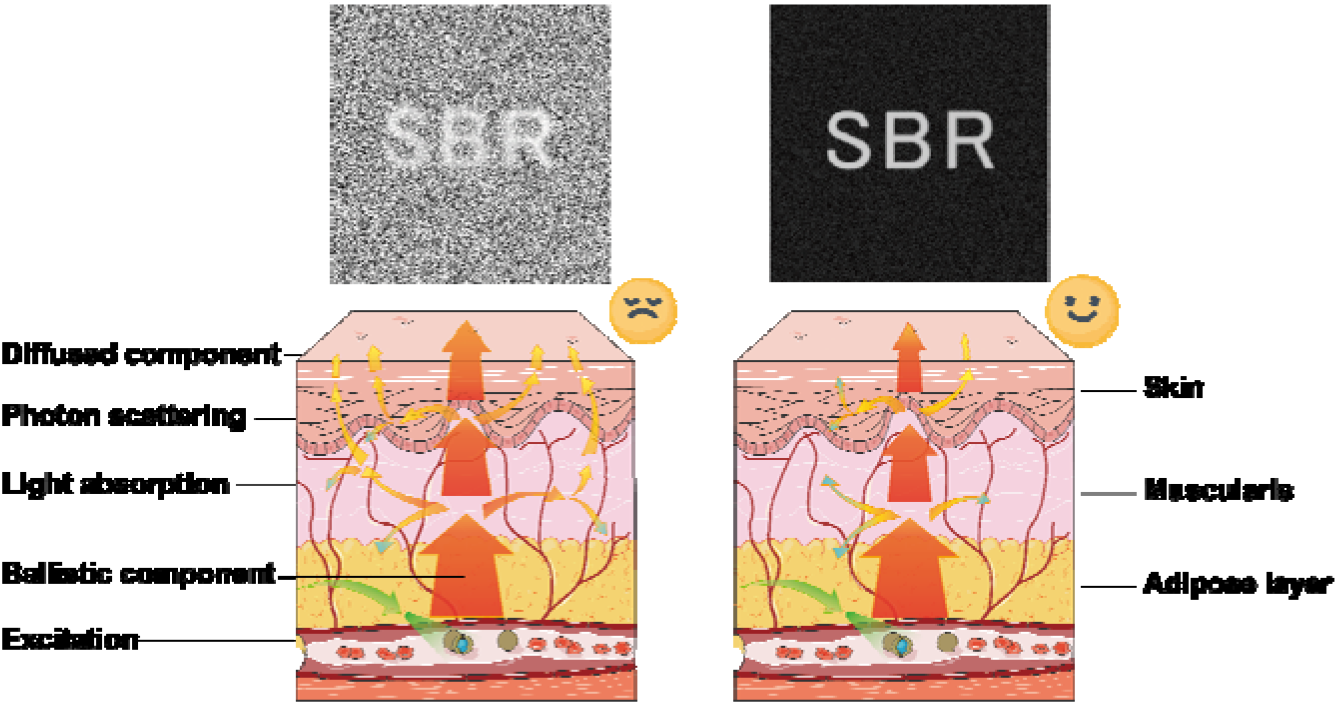
The light propagation in the bio-tissue far away from (left) and around (right) the light absorption peak.

**Fig. 1.**
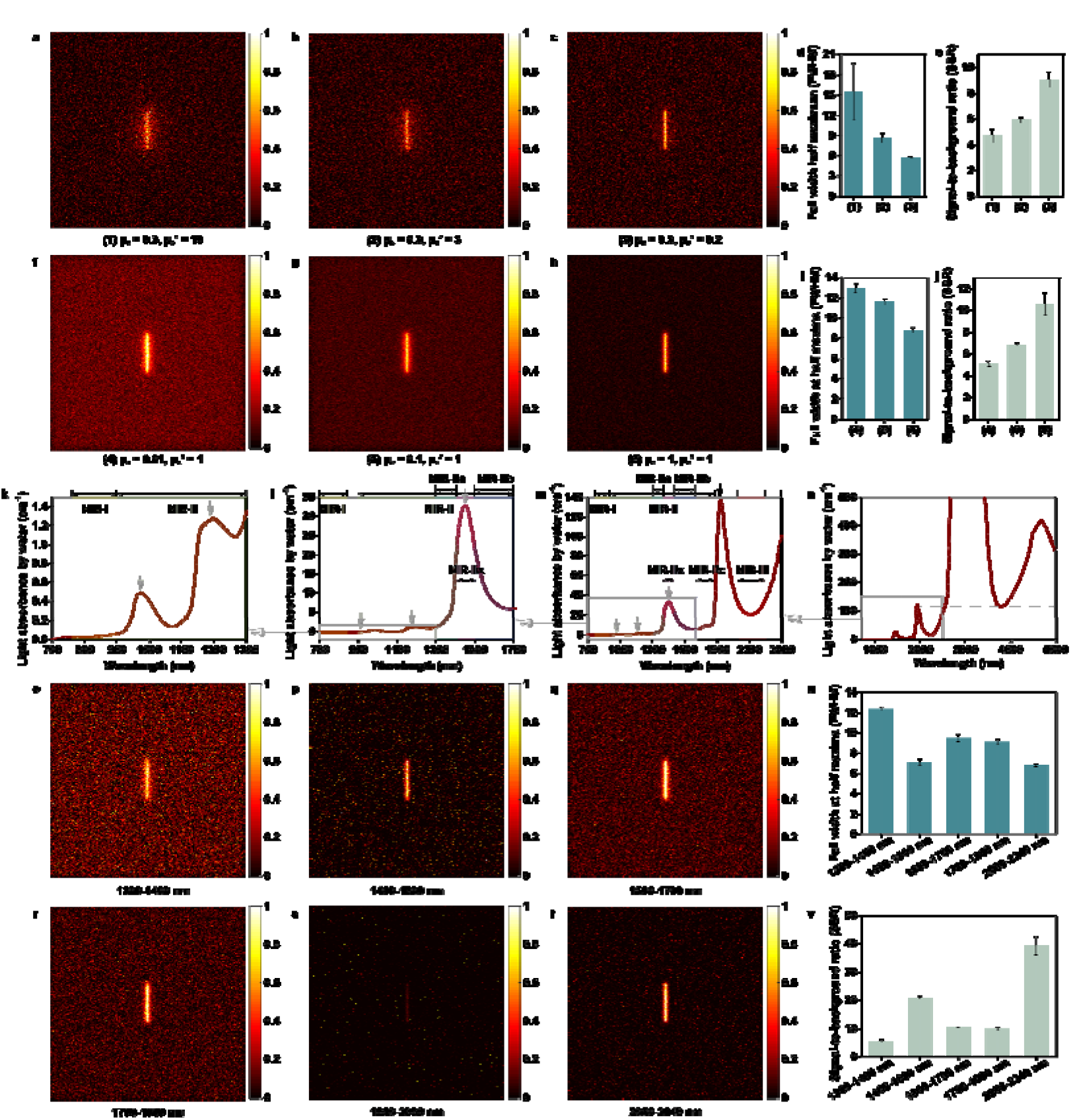
The simulation results of near-infrared bio-tissue imaging via Monte Carlo method. Images of a line source through a bio-tissue of 1 mm with scattering anisotropy factor (g) of 0.9, absorption coefficient (μ_a_) of 0.3 mm^−1^ and varying reduced scattering coefficient (μ_s_') **(a)** μ_s’ =_ 10 mm^−1^, **(b)** μ_s’ =_ 3 mm^−1^ and **(c)** μ_s’ =_ 0.2 mm^−1^. The **(d)** full width at half maxima (FWHM) and **(e)** signal-to-background ratio (SBR) analyses of the samples in fig. 1a-c. Images of a line source through a bio-tissue of 1 mm with g of 0.9, μ_s’_ of 1 mm^−1^ and varying absorption coefficient **(f)** μ_a_ = 0.01 mm^−1^, **(g)** μ_a_ = 0.1 mm^−1^ and **(h)** μ_a_ = 1 mm^−1^. The **(i)** FWHM and **(j)** SBR analyses of the samples in Fig. 1f-h. The light absorption spectra of water within **(k)** 700-1300 nm, **(l)** 700-1700 nm, **(m)** 700-2500 nm and **(n)** 900-5000 nm. The gray arrows pointed the absorption peaks and the gray dashed line in Fig. 1n represented the absorbance at 1930 nm. Images of a line source in **(o)** 1300-1400 nm, **(p)** 1400-1500 nm, **(q)** 1500-1700 nm, **(r)** 1700-1880 nm, **(s)** 1880-2080 nm and **(t)** 2080-2340 nm. The **(u)** FWHM and **(v)** SBR analyses of the samples in Fig. 1o-t.

The light absorption spectra within 700-5000 nm of water, which is the most important component of organisms, were shown in Fig. 1k-n.^40,41^ As is widely known to all, the 360-760 nm is defined as visual region, the NIR region thus starts with 760 nm. Traditional bio-imaging window is usually located in NIR-I region, which is ranging from 760 nm to 900 nm^42,43^. The light absorption by water substantially improves beyond 900 nm (Fig. 1k), compared with NIR-I window. The gray arrows pointed the absorption peaks at ~980 nm, ~1200 nm, ~1450 nm and ~1930 nm, where the spectral regions around the peaks would improve the imaging quality in principle. Because of the absorption peak at ~980 nm, 900-1000 nm should not be excluded from the NIR-II window for bio-imaging. The imaging in 1400-1500 nm has not long been recognized, but the high light absorption within this band, which is called as NIR-IIx region here, is no longer the barrier in NIR-II region, as long as the fluorescent probes possess enough brightness to resist the attenuation by water. At present, the photoresponse of the classic InGaAs detector limited the optical imaging beyond 1700 nm, thus NIR-II window was defined as no more than 1700 nm. Because of the similar absorption and scattering properties, we believed that 1700-1880 nm possessed comparable imaging quality with the NIR-IIb imaging and defined 1700-1880 nm as the NIR-IIc region. It could be calculated that the peak absorption intensity at ~1930 nm is near e^100^ times higher than the peak at ~1450 nm for every 1 cm of transmission, thus the useful signals with the wavelengths near 1930 nm would be almost impossible to detect in deep tissue. Hence, the NIR-II window was perfected as 900-1880 nm. Over the absorption “mountain” peaking at ~1930 nm, the region of 2080-2340 nm, which was newly considered as the third near infrared (NIR-III) region here, becomes the last high-potential bio-window in general since the water absorption of light beyond 2340 nm keeps stubbornly high (Fig. 1n).

In order to verify the above hypotheses, we then simulated the imaging in 1300-1400 nm (NIR-IIa), 1400-1500 nm (NIR-IIx), 1500-1700 nm (NIR-IIb), 1700-1880 nm (NIR-IIc), 1880-2080 nm and 2080-2340 nm (NIR-III) window (Fig. 1o-t), taking the absorption spectrum of water and the scattering property of skin into consideration. The average of multiple simulated images (displayed in Fig. S2) within one NIR region were considered as the equivalent result of the window. As shown in Fig. 1u-v, except for the extremely intense depletion in 1880-2080 nm (Fig. 1s), rising light absorption and falling photon scattering both made positive contributions to the precise imaging. Interestingly, the NIR-IIx imaging showed the best FWHM and SBR in the whole NIR-II region. Compared with the NIR-IIb region, the production of scattering photons was less but the depletion of scattering photons (determined by absorption) dropped at the same time in NIR-IIc region. Altogether, the NIR-IIc imaging also performed well with good definition and high SBR. Furthermore, since the subcutaneous adipose absorbs more light in NIR-IIc region, the NIR-IIc imaging holds greater potential in obese organisms.^44^ Significantly, the newly defined NIR-III imaging possessed greatest potential for bioimaging owing to the large but proper absorbance and the exceedingly low photon scattering, according to the results from Monte Carlo simulation. We believed the NIR-IIc and NIR-III imaging would be achieved sooner or later with the rapid development and further popularization of NIR detectors as well as the emergence of bright agents.

### 900-1000 nm should be included in the second near-infrared window

The second near-infrared region has long been defined as 1000-1700 nm in academic world.^45^ Meanwhile, many InGaAs-based products enable imaging from 0.9 μm to 1.7 μm and this range has also been a typical band of shortwave infrared in industrial field. The boundary of the two similar regions has been ambiguous. In particular, 900-1000 nm seems to have not been recognized as part of NIR-II region by most research groups. According to the advantages of moderate light absorption mentioned above, 900-1000 nm might be a promising imaging window considering that there was an absorption peak at ~980 nm.

To examine the imaging window in NIR-II region, the PbS QDs were developed, whose brilliant fluorescence intensity and stability would facilitate the long-time monitoring and tracking *in vivo*.^46,47^ Besides, the broad excitation with narrow and asymmetry emission of PbS QDs could enable multiplex imaging *in vivo*,^48,49^ compared with other NIR-II emitting fluorophores. Firstly, the PbS/CdS QDs emitting at ~1100 nm were developed, which were named as 1100-PbS/CdS QDs. The absorption and emission spectra of 1100-PbS/CdS QDs in tetrachloroethylene were displayed in Fig. S3. As shown in Fig. 2a & b, the PEGylated 1100-PbS/CdS QDs showed bight emission in NIR-II region and the integrated emission intensities of the 1100-PbS/CdS QDs in 900-1000 nm were weaker than those in 1000-1100 nm. It could be seen that though the 900-1000 nm window possessed higher photon scattering, the light absorption by water in that region was also stronger, compared with the band of 1000-1100 nm (Fig. 2c). The transmission electron microscopic (TEM) image (Fig. S4a) and the dynamic light scattering (DLS) result (Fig. S4b) identified the uniformly spherical QDs with a mean hydrodynamic diameter of 29.9 nm. The zeta potential was measured as −2.0 mV (Fig. S4c).

**Fig. 2.**
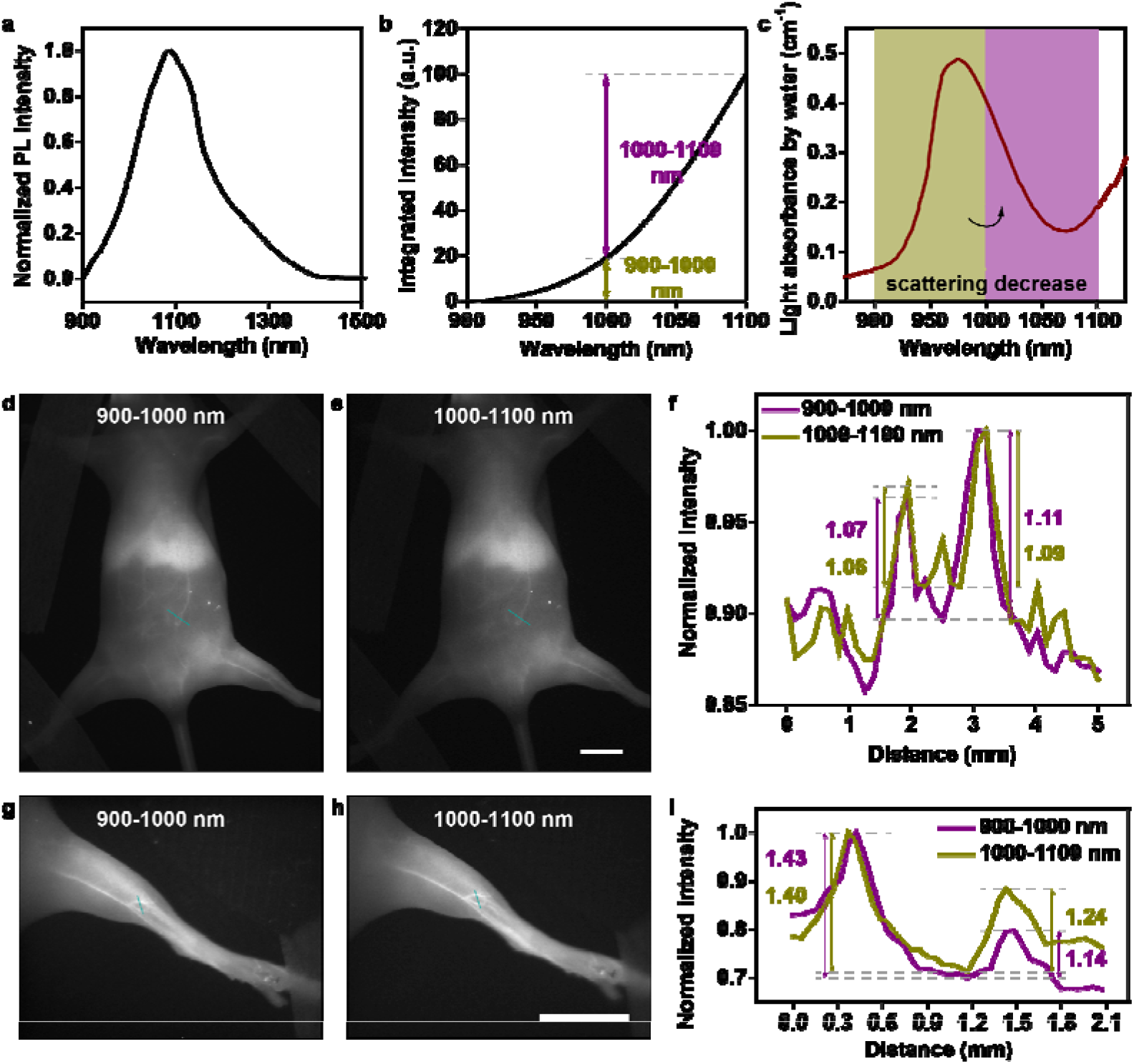
Comparison of *in vivo* fluorescence imaging in 900-1000 nm and 1000-1100 nm. **(a)** The emission spectrum of PEGylated 1100-PbS/CdS QDs in water. **(b)** The integrated emission intensity of PEGylated 1100-PbS/CdS QDs in water. **(c)** The light absorbance of water within 800-1200 nm. The whole-body imaging in **(d)** 900-1000 nm and **(e)** 1000-1100 nm. **(f)** Cross-sectional fluorescence intensity profiles along the indigo lines of the blood vessel in Fig. 2d & e. The numbers show the SBRs. The hind limb imaging in **(g)** 900-1000 nm and **(h)** 1000-1100 nm. **(i)** Cross-sectional fluorescence intensity profiles along the indigo lines of the blood vessel in Fig. 2g & h. The numbers show the SBRs. Scale bars in (h) & (k), 10 mm.

*In vivo* fluorescence imaging was then conducted under the excitation of 793 nm CW laser with a power density of ~2 mW cm^−2^. As shown in Fig. 2d & e, despite the dominant intensity proportion within 1000-1100 nm of the PEGylated 1100-PbS/CdS QDs, the whole-body imaging in 900-1000 nm is not significantly worse than that in 1000-1100 nm post intravenous injection (1 mg mL^−1^, 200 μL). In contrast, the quantitative results in Fig. 2f even gave us strong evidence about the better SBRs of 1.07 and 1.11 in 900-1000-nm imaging, while the SBRs of the same vessels were 1.06 and 1.09 in 1000-1100-nm imaging. By adjusting the field of view, the whole hind limb could be clearly presented (Fig. 2g & h; Power density of the excitation, ~12 mW cm^−2^), giving the further comparison of imaging quality between 900-1000 nm and 1000-1100 nm and the contrasts in the images were demonstrated to be level-pegging, which is shown in Fig. 2i. That is to say, though the photon scattering decreases with the increase of wavelength, the strong absorption at ~980 nm could actually suppress the image background. Besides, the distinct advantages of imaging in 900-1000 nm was verified by other groups compared with the imaging in 700-900 nm.^50^ In view of the same detection mechanism and decent imaging performance between the spectral region of 900-1000 nm and 1000-1100 nm, the former should not be excluded from the NIR-II imaging window.

### The rising light absorption by water from ~1300 nm “turns on” the promising stage for NIR-II fluorescence imaging

The PbS/CdS CSQDs emitting at ~1300 nm were next synthesized for the comparison of the imaging in 1100-1300 nm and 1300-1500 nm, whose absorption and emission spectra were shown in Fig. S5. After PEGylation, the 1300-PbS/CdS QDs were spherical with a hydrodynamic size of 35.2 nm from the TEM (Fig. S6a) and DLS (Fig. S6b) results and the zeta potential was measured as −2.8 mV (Fig. S6c). The normalized and integrated emission spectra of the PEGylated 1300-PbS/CdS QDs were presented in Fig. 3a-b and the calculated fluorescence intensities in 1300-1500 nm were marginally lower than those in 1100-1300 nm, which might be blamed on the light absorption by water. It was apparent from Fig. 3c that the shifting from 1100-1300 nm to 1300-1500 nm not only decreased the photon scattering, but also noteworthy heightened the light absorption, which would lead to a remarkable improvement on the background suppression.

**Fig. 3.**
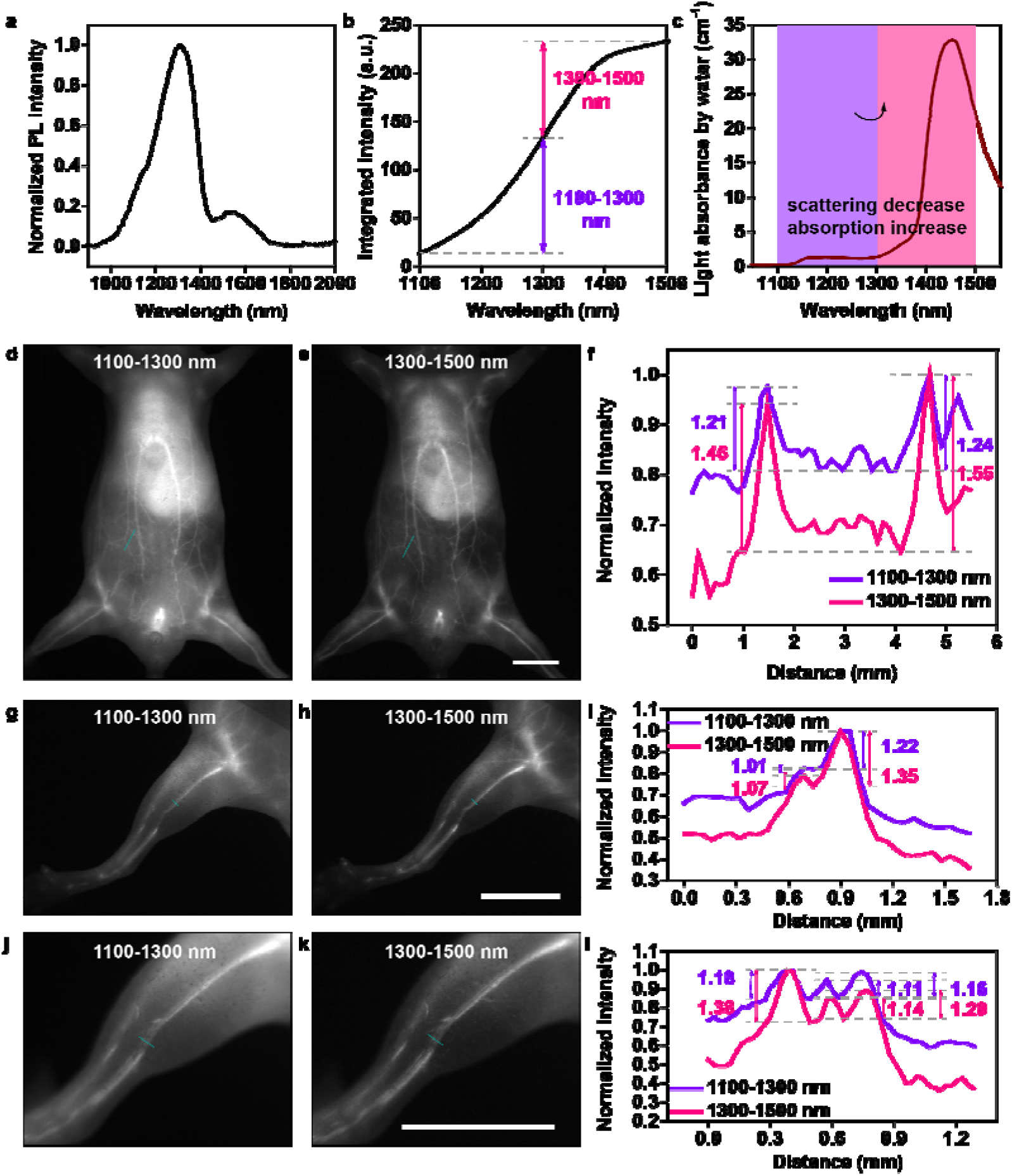
Comparison of *in vivo* fluorescence imaging in 1100-1300 nm and 1300-1500 nm. **(a)** The emission spectrum of PEGylated 1300-PbS/CdS QDs in water. **(b)** The integrated emission intensity of PEGylated 1300-PbS/CdS QDs in water. **(c)** The light absorbance of water within 1000-1600 nm. The whole-body imaging in **(d)** 1100-1300 nm and **(e)** 1300-1500 nm. **(f)** Cross-sectional fluorescence intensity profiles along the indigo lines of the blood vessel in Fig. 3d & e. The numbers show the SBRs. The hind limb imaging in **(g)** 1100-1300 nm and **(h)** 1300-1500 nm. **(i)** Cross-sectional fluorescence intensity profiles along the indigo lines of the blood vessel in Fig. 3g & h. The numbers show the SBRs. The hind limb imaging at a larger magnification in **(j)** 1100-1300 nm and **(k)** 1300-1500 nm. **(l)** Cross-sectional fluorescence intensity profiles along the indigo lines of the blood vessel in Fig. 3j & k. The numbers show the SBRs. Scale bar, 10 mm.

After intravenous injection (1 mg mL^−1^, 200 μL), the whole-body vessel imaging could be conducted under the excitation of 793 nm CW laser (Power density of the excitation in Fig. 3d, ~3 mW cm^−2^; Power density of the excitation in Fig. 3e, ~30 mW cm^−2^). As displayed in Fig. 3d & e, the image background decreased significantly when shifting the spectral window from 1100-1300 nm to 1300-1500 nm. After calculation, the SBRs showed uplifts over 20 per cent (Fig. 3f). The hind limb vessel imaging was then performed in the two bands (Power density of the excitation in Fig. 3g, ~15 mW cm^−2^; Power density of the excitation in Fig. 3h, ~37 mW cm^−2^) and the high-contrast advantage in 1300-1500-nm imaging was demonstrated once again (Fig. 3g-i). When further enlarging the magnification, the adjacent three fine vascular structures could be still distinguishable with low background in 1300-1500 nm (Fig. 32j-l; Power density of the excitation in Fig. 3j-k, ~60 mW cm^−2^) and each capillary exhibited the better SBRs of 1.38, 1.14 and 1.20 than those in 1100-1300 nm (1.18, 1.11 and 1.16). Typically, imaging window beyond 1300 nm, which was called as NIR-IIa region, has been already verified eminently suitable for fluorescence bioimaging throughout the whole NIR-II region. It could be seen that the 200-nm red shift induced scattering depression and the surge in the absorption in especial contributed to the background attenuation. Then we injected the 1100-PbS/CdS QDs into another mouse, and conducted the 900-1100-nm and 1100-1300-nm imaging. However, Fig S7 showed that just the 200-nm red shift from 900-1100 nm to 1100-1300 nm did not similarly improve the imaging quality. It seemed that the substantial growth of light absorption but not the decrescent photon scattering governed the start of promising stage for NIR-II fluorescence imaging.

### The optimum NIR-II imaging window locates around the intense absorption at ~1450 nm

The intense absorption by water peak locating in the NIR-II region has been eschewed for its strong signal loss and potential thermal damage. To objectively evaluate the fluorescence imaging with collection around 1450 nm, the 1450-PbS/CdS QDs were synthesized whose absorption and emission spectra could be found in Fig. S8. Plainly, the TEM (Fig. S9a) and DLS (Fig. S9b) results of PEGylated 1450-PbS/CdS QDs demonstrated a uniform spherical shape with an average hydrodynamic size of 35.2 nm and the zeta potential of −3.8 mV (Fig. S9c). The hydrogen bond and the O-H covalently bond in water make the absorption spectrum present multiple characteristic peaks in infrared region. When the protium in water is replaced by deuterium, the molecular weights increase, and the characteristic absorption peak wavelength at ~1450 nm is red shifted to ~1970 nm. The normalized photoluminescence spectra of PEGylated 1450-PbS/CdS QDs in hydrogen oxide (water) and deuterium oxide (heavy water) were exhibited in Fig. 4a. It was apparent that the intense absorption at ~1450 nm by water induced a large depression in the fluorescence spectrum, while the spectrum of the QDs in heavy water, where the corresponding absorption peak was red shifted, restored the original emission characteristics. After integration, it was noticeable that the emission in water within 1425-1475 nm was depleted seriously (Fig. 4b). Fig. 4c directly presented the light absorption within 1300-1800 nm. The 1300-1400 nm and 1500-1700 nm were called as NIR-IIa and NIR-IIb region, respectively, where the fluorescence imaging was proved with excellent quality. We now define the neglected region of 1400-1500 nm as NIR-IIx window, which has long been considered unsuitable for imaging owing to the highest absorbance at ~1450 nm in NIR-II region.

**Fig. 4.**
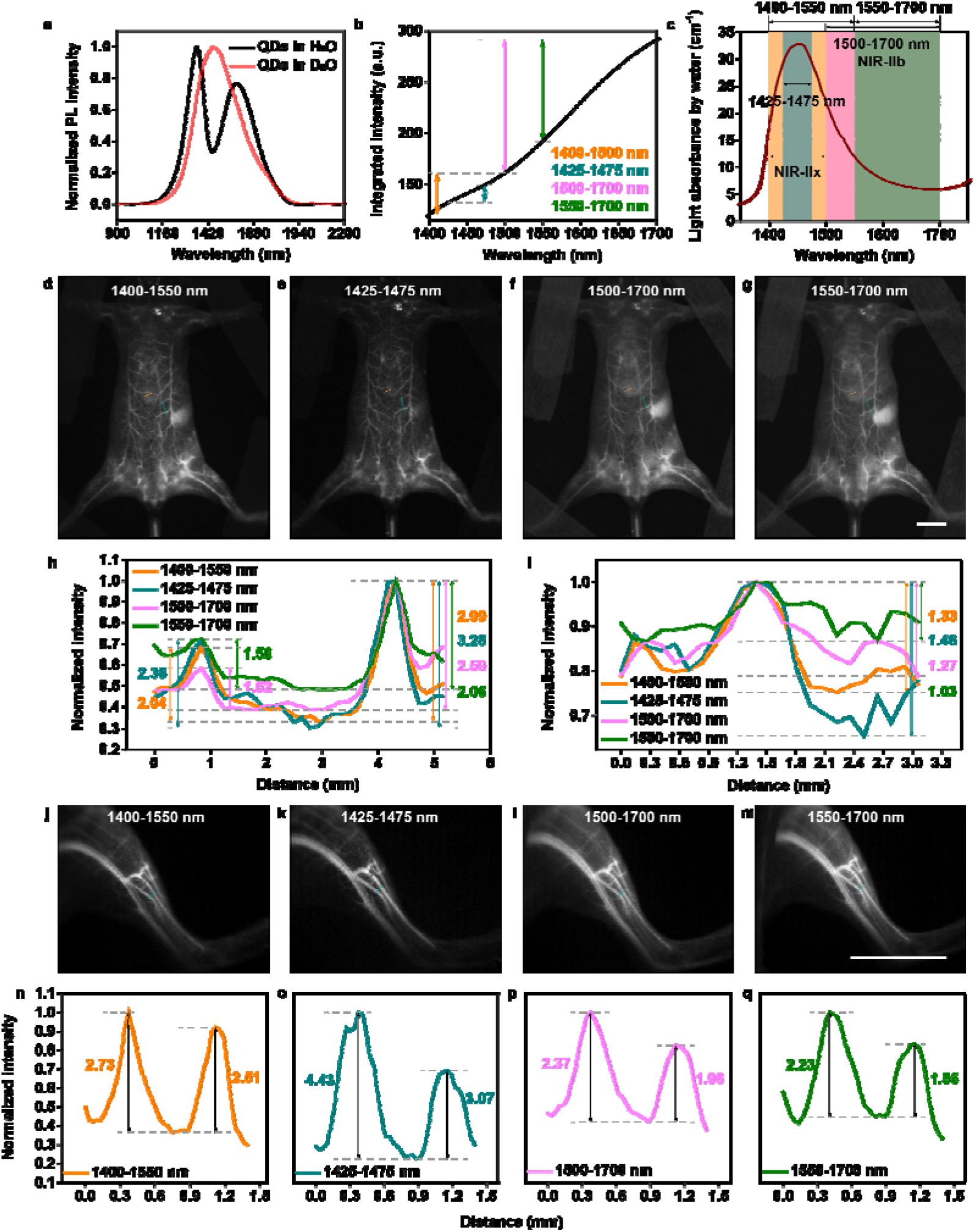
Comparison of *in vivo* fluorescence imaging in 1400-1550 nm, 1425-1475 nm, 1500-1700 nm and 1550-1700 nm. **(a)** The emission spectra of PEGylated 1450-PbS/CdS QDs in water and heavy water. **(b)** The integrated emission intensity of PEGylated 1450-PbS/CdS QDs in water. **(c)** The light absorbance of water within 1300-1800 nm. The whole-body imaging in **(d)** 1400-1550 nm, **(e)** 1425-1475 nm, **(f)** 1500-1700 nm and **(g)**1550-1700 nm. Cross-sectional fluorescence intensity profiles along **(h)** the indigo lines and **(i)** brown lines of the blood vessel in Fig. 4d-g. The numbers show the SBRs. The hind limb imaging in **(j)** 1400-1550 nm, **(k)** 1425-1475 nm, **(l)** 1500-1700 nm and **(m)**1550-1700 nm. **(n-q)** Cross-sectional fluorescence intensity profiles along the indigo lines of the blood vessel in Fig. 4j-m. The numbers show the SBRs. Scale bar, 10 mm.

Under the excitation of 793 nm CW laser, the whole-body vessel imaging was conducted in 1400-1500 nm, 1425-1475 nm, 1500-1700 nm and 1550-1700 nm. It is worth noting that the 1425-1475 nm possesses a bandwidth of 50 nm around the absorption peak at ~1450 nm, which highlights the contribution of absorption by water; The band of 1400-1500 nm broadens the detection around 1450 nm with a bandwidth of 100 nm, which is the previously avoided region and defined as NIR-IIx region here; The 1500-1700 nm is the classic NIR-IIb region with existing excellent performance; The 1550-1700 nm is the chosen region with the lowest photon scattering compared with the above three bands. As shown in Fig. 4d-g (Power density of the excitation in Fig. 4d & e, ~40 mW cm^−2^; Power density of the excitation in Fig. 4f&g, ~20 mW cm^−2^), the vessel imaging in 1425-1475 nm possessed the lowest imaging background. It is interesting to note that increasing the imaging wavelength did not make the performance as better as expected, since the absorption peak located at ~1450 nm. Comparing the imaging in the four regions, the photon scattering in 1550-1700 nm is theoretically lowest but the region with high expectations gave the worst image contrast. Meanwhile, the imaging in 1425-1475 nm provided the best SBRs of 2.36 and 3.28 and with the imaging region shifted farther away from the absorption peak, the SBRs became worse (2.04 and 2.99 in 1400-1550 nm, 1.52 and 2.59 in 1500-1700 nm, 1.50 and 2.06 in 1550-1700 nm), which is shown in Fig. 4h. On account of the accumulation of the QDs in liver and spleen, the strong background blurred the vessels above the two bright organs. However, as shown in Fig. 4i, the SBRs were calculated as 1.33, 1.46, 1.27 and 1.03 respectively in 1400-1500-nm, 1425-1475-nm, 1500-1700-nm and 1550-1700-nm images, which revealed that the absorption induced background attenuation could improve the definition effectively. The hind limb imaging with larger magnification was also conducted in the four regions. It could be seen in Fig. 4j-m that, the closer to the imaging window is to the peak absorption, the lower the imaging background (Power density of the excitation, ~40 mW cm^−2^). As shown in Fig. 4n-q, the SBRs of the two tiny vessels were measured as 2.73 and 2.51 in 1400-1500 nm, 4.43 and 3.07 in 1425-1475 nm, 2.37 and 1.96 in 1500-1700 nm, 2.23 and 1.86 in 1550-1700 nm, which further confirmed the positive contribution of the absorption. NIR-IIb fluorescence imaging has long been regarded as the most promising fluorescence imaging technique due to the suppressed photon scattering until then, but our results now proved the greater contribution of rising absorption than the decrescent scattering and the NIR-IIx fluorescence imaging proposed in this work owned optimum performance, even exceeding the NIR-IIb fluorescence imaging.

Deep-penetration fluorescence hysterography gives a promising diagnostic method in uterine anomalies and intrauterine lesions with multiple advantages of non-invasive property, high spatial resolution, and no ionizing radiation. Besides, bladder is also a hollow organ in urinary system, that stores and controls urine. The bladder fluorescence imaging benefits precisely monitoring the volume variation which may be related to lower urinary tract symptoms including storage disorder. Fig. 5 showed the results of hollow organ imaging and the definitions of imaging in different imaging window were further compared. After uterine perfusion (1 mg mL^−1^, 200 μL), the bright and specific uterine structure was presented. In uterine imaging (Fig. 5a-c; Power density of the excitation, ~30 mW cm^−2^), the skin, fatty, muscularis and the organs above, such as gut and bladder, were a brake on the photon propagation, leading to a diffuse structural boundary. The image in 1425-1475 nm (Fig. 5b) gave us the sharpest outline in comparison. The quantitative intensities and the results of Gaussian fitting were displayed in Fig. 5d-f and the FWHMs of the same site were 1.97 mm, 1.70 mm, and 2.05 mm, respectively. Obviously, the highly weakening of background induced by photon scattering could be concluded that not only enormously increase the image contrast (Fig. 4), but also improve the spatial resolution of the details deeply buried in the bio-tissue (Fig. 5). Moreover, the bladder was labeled via intravesical perfusion (1 mg mL^−1^, ~20 μL). As shown in Fig. 5g-m (Power density of the excitation, ~10 mW cm^−2^), the imaging quality did not improve with the increase of imaging wavelength, which was not consistent with the general understanding before. Through the skin and muscles, the bladder image showed the clearest contour in 1425-1475 nm with a narrowest fitted diameter of 4.24 mm, while the FWHMs of the bladders in 1400-1550-nm and 1500-1700-nm (NIR-IIb) imaging were 4.62 mm and 4.95 mm. In view of this, the NIR-IIx fluorescence imaging deciphered the deep details *in vivo* precisely, which held the strong potential to advance the medical imaging in clinic.

**Fig. 5.**
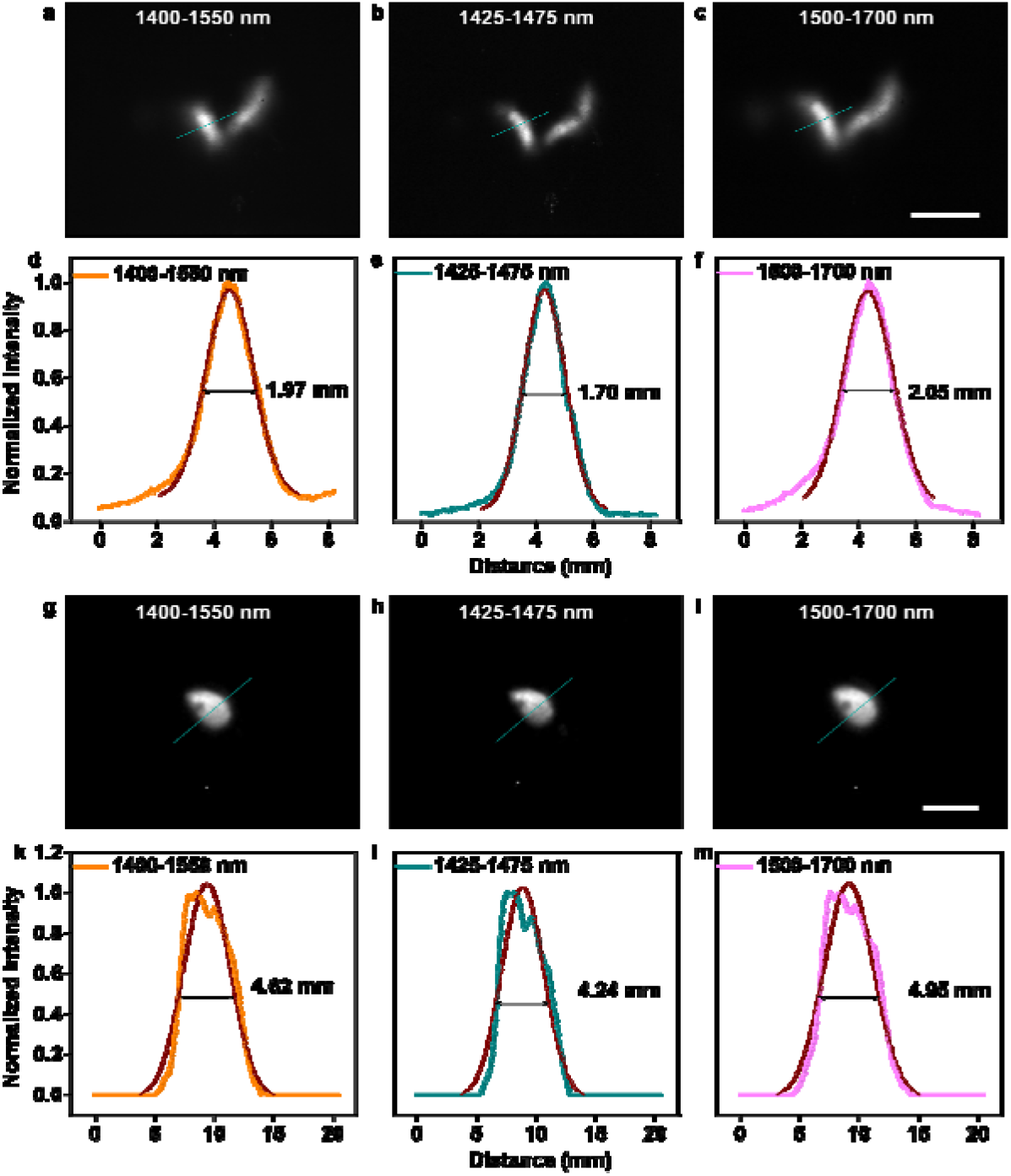
Functional *in vivo* fluorescence imaging in 1400-1550 nm, 1425-1475 nm and 1500-1700 nm. The uterine imaging in **(a)** 1400-1550 nm, **(b)** 1425-1475 nm and 1500-1700 nm. **(d-f)** The fluorescence intensity analyses of the right uterus (the indigo lines) in Fig. 5a-c. The numbers show the FWHMs. The bladder imaging in **(g)** 1400-1550 nm, **(h)** 1425-1475 nm and **(i)** 1500-1700 nm. **(k-m)** The fluorescence intensity analyses of the bladder (the indigo lines) in Fig. 5g-i. The numbers show the FWHMs. Scale bar, 10 mm.

### Large-depth fluorescence tomography via wide-field microscopy around NIR-IIx region

User-friendly fluorescence wide-field microscopy, as a classical technique, was often utilized for cell or tissue slice imaging. In recent years, the imaging window of wide-field microscopy has been shifted into the near-infrared region to shrink the photon scattering and visualize through the deep tissue *ex* or *in vivo*. Nowadays, NIR-II fluorescence wide-field microscopy has succeeded in penetrating ~800 μm below the skull in the brain. However, despite the large imaging depth, the scattering photons and the signal photons from outside the focal plane induced background kept the details hidden beneath a veil of “mist”. With the excellent SBR and resolution of the imaging around the peak absorption wavelength shown above, wide-field microscopy around the NIR-IIx region was believed to possess excellent performance without complex excitation and collection modes.

The 5× imaging was conduct with a scan lens (LSM03, Thorlabs) equipped. As shown in Fig. 6a-d, it could be seen that the light absorption by water greatly restrained the background. The quantitative results were presented in Fig. 6e-h, and the SBRs were 1.80 and 2.40 in 1400-1550-nm image, 2.84 and 4.47 in 1425-1475-nm image, 1.62 and 2.18 in 1500-1700-nm image, 1.58 and 1.92 in 1550-1700-nm image, respectively. Inevitably, the strong light absorption would make losses of useful signals meanwhile, in spite of remarkable contrast improvement. In order to efficiently reduce the signal loss in deep exploring, 1400-1550 nm was then determined as the imaging region for microscopy with larger magnification. The 25× cerebral vasculature images at the depth of 150 μm and 650 μm below the skull were displayed in Fig. 6i-l. The three capillaries in 1400-1500-nm image all showed higher SBRs of 2.69, 4.11 and 1.77 than those in NIR-IIb image (2.04, 2.89 and 1.58) at 150 μm. Besides, the 650-μm-depth vasculum imaged in 1400-1500 nm possessed the SBR of 4.75 while the NIR-IIb image gave the SBR of 4.35 when measuring the same vessel. As shown in Fig. 6q, the images of cortex vessels kept extremely low background within ~500 μm in the brain. At ~900 μm below the skull, the vessel network was still clearly visible and the sharp and tiny capillary with a FWHM of just 4.1 μm could be easily distinguished. Beyond 900 μm, the potential white matter might become the obstacles for further visualizing and the image details started to become sparse. At about ~1.3 mm below the skull, there still existed recognizable vascular, which represented the largest imaging depth of *in vivo* NIR-II fluorescence microscopy in the mice brain up to now.

**Fig. 6.**
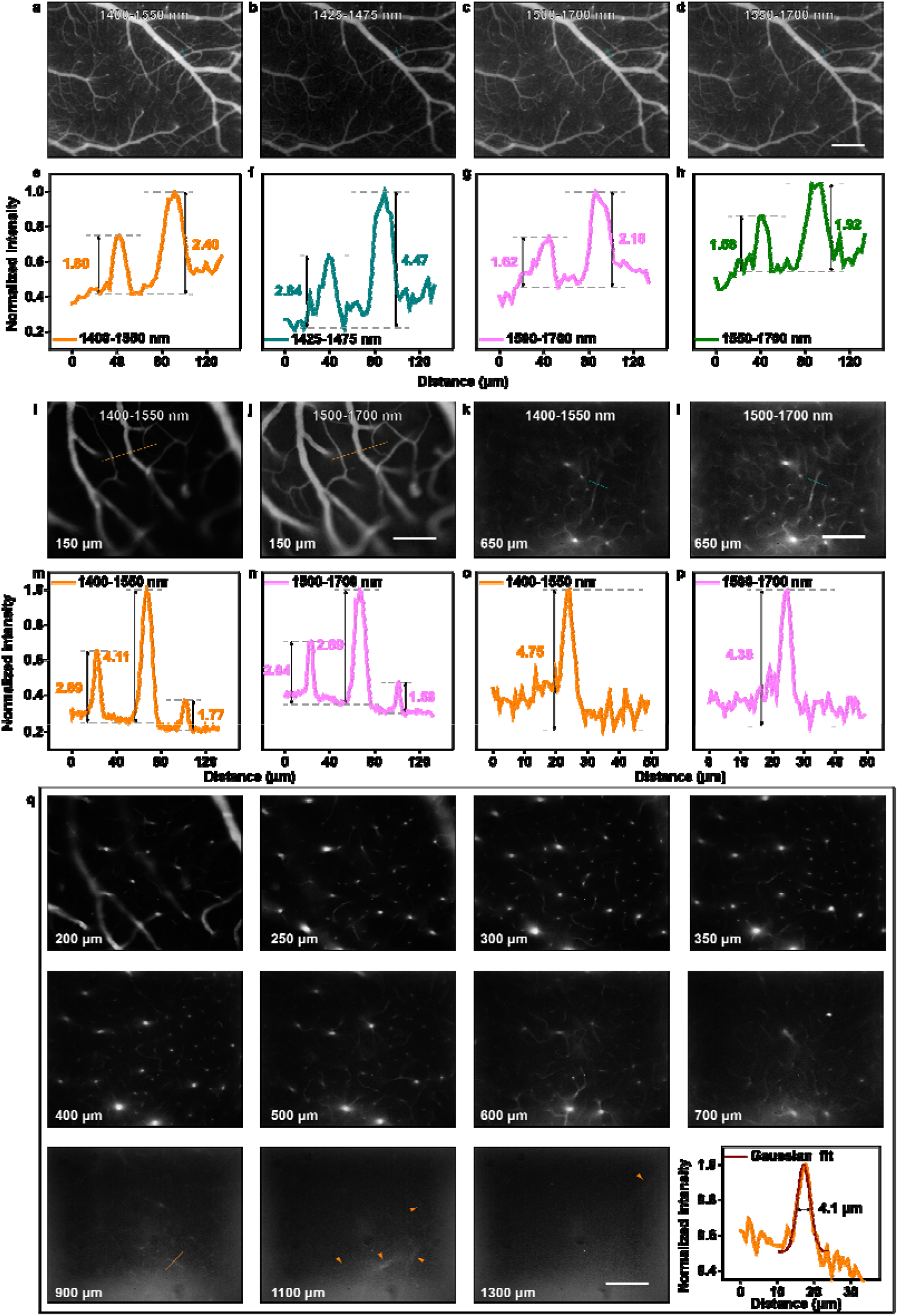
*In vivo* fluorescence wide-field microscopic imaging beyond 1400 nm. The 5× cerebral vasculature imaging in **(a)** 1400-1550 nm, **(b)** 1425-1475 nm, **(c)** 1500-1700 nm and **(d)**1550-1700 nm. Scale bar, 300 μm. **(e-h)** Cross-sectional fluorescence intensity profiles along the indigo lines of the blood vessel in Fig. 6a-d. The numbers show the SBRs. The 25× cerebral vasculature imaging at depth of 150 μm in **(i)** 1400-1550 nm and **(j)** 1550-1700 nm. Scale bar, 100 μm. The 25× cerebral vasculature imaging at depth of 650 μm in **(k)** 1400-1550 nm and **(l)** 1550-1700 nm. Scale bar, 100 μm. Cross-sectional fluorescence intensity profiles along **(m-n)** the indigo lines in Fig. 6i-j and **(o-p)** the brown lines in Fig. 6k-l. The numbers show the SBRs. **(q)** 25× cerebral vasculature imaging in 1400-1550 nm at various depth below the skull and the fluorescence intensity analysis of the blood vessel at the depth of 900 μm (the brown line). The brown arrows show the deep and tiny capillaries. Scale bar, 100 μm.

### Off-peak NIR-II fluorescence imaging goes best with 1400 nm long-pass (NIR-IIx+NIR-IIb) collection

Nowadays, many efforts have been made for design and synthesis of imaging probes with both long emission wavelength and strong emission, especially for organic dyes. The recent researches on the optimization of off-peak NIR-II fluorescence imaging has also expedited the development of bright NIR-II emissive agents. Owing to the considerable emission and the broad band, some NIR emissive dyes were successfully utilized for NIR-IIb (beyond 1500 nm) fluorescence imaging, which was long-term routinely described as the best NIR-II imaging window resulting from the minimized photon scattering. As a matter of fact, the poorer performance in 1550-1700 nm, compared to the imaging in 1500-1700 nm (NIR-IIb), has made it clear that simply reducing the scattering could not promote the imaging quality (Fig. 4i-j). Thus, we believed that the improvement of NIR-IIb window was attributed in large part to the light absorption, in addition, 1400-1500 nm region, which entirely included the huge absorption peak at ~1450 nm, shouldn’t be filtered in off-peak fluorescence imaging theoretically.

As a representative NIR-II dye reported in our previous work,^7^ IDSe-IC2F was chosen to explore the optimum imaging window for off-peak NIR-II fluorescence imaging. The molecular structure was shown in Fig. 7a in which 7,7′◻(4,4,9,9◻tetrakis(4◻hexylphenyl)◻4,9◻dihydro◻s◻indaceno[1,2◻b:5,6◻b′]bi s(selenophene)◻2,7◻diyl)bis(2,3◻dihydrothieno[3,4◻b][1,4]dioxine◻5◻carbaldehy de) and 2◻(5,6◻difluoro◻3◻oxo◻2,3◻dihydro◻1H◻inden◻1◻ylidene)malononitrile served as the electron◻donating and ◻withdrawing moieties respectively with a π◻bridge of ethylene dioxothiophene (EDOT) unit. To improve the solubility, the IDSe-IC2F molecules were encapsulated into nanoparticles via PEGylation (Fig. 7b). The hydrated IDSe-IC2F NPs acted as the typical NIR-II fluorophores with absorption peaks in NIR-I region and the emission peak in NIR-II region. Their absorption and emission spectra were given in Fig. 7c-d and the integrated fluorescence intensities in Fig. 7e showed the tail emission.

After intravenous injection of the IDSe-IC2F NPs (1 mg/mL, 200 μL), the whole-body imaging in mice was conducted under the excitation of 793 nm CW laser (Fig. 7f-l). The images in 900/1000/1100/1200/1300/1400/1500-1700 nm showed varying degrees of absorption and scattering in tissue (The power densities of the excitation were ~1 mW cm^−2^, ~2 mW cm^−2^, ~30 mW cm^−2^, ~55 mW cm^−2^, ~75 mW cm^−2^, ~110 mW cm^−2^ and ~110 mW cm^−2^ in Fig. 7f, 7g, 7h, 7i, 7j, 7k and 7l, respectively.). After calculation, the selected three vessels (indigo lines) showed the highest SBRs of 1.72, 2.07 and 1.75 in 1400-nm-LP image, while those in 900-nm-LP, 1000-nm-LP, 1100-nm-LP, 1200-nm- LP, 1300-nm-LP, 1500-nm-LP images were 1.33, 1.33 and 1.21, 1.36, 1.30 and 1.13, 1.42, 1.47 and 1.25, 1.57, 1.69 and 1.37, 1.64, 1.77 and 1.35, 1.61, 1.87 and 1.40, respectively. Besides, the hind limb imaging was further carried out and the background suppression near the absorption peak by water was obvious in Fig. 7n-t (The power densities of the excitation were ~1.5 mW cm^−2^, ~2 mW cm^−2^, ~40 mW cm^−2^, ~70 mW cm^−2^, ~100 mW cm^−2^, ~150 mW cm^−2^ and ~150 mW cm^−2^ in Fig. 7n, 7o, 7p, 7q, 7r, 7s and 7t, respectively.). The SBRs of the selected two vessels reached the maximum of 8.81 and 11.68 when imaging in 1400-1700 nm (Fig. 7u). It could be concluded that the performance of NIR-IIx+NIR-IIb (1400-1700 nm) imaging surpassed the NIR-IIb imaging since the NIR-IIx signals did positive contribution with only slight background added. Moreover, the 900-nm-LP fluorescence imaging did no worse than the 1000-nm-LP fluorescence imaging, which once again demonstrated that 900-1000 nm should not be excluded from the NIR-II region. In the images at 2 hours post injection, the liver was lit up (Fig. 7v-x) since substantial IDSe-IC2F NPs in blood accumulated in the liver (The power densities of the excitation were ~75 mW cm^−2^, ~110 mW cm^−2^ and ~110 mW cm^−2^ in Fig. 7v, 7w and 7x, respectively.). As shown in Fig. 7y, the NIR-IIx+NIR-IIb fluorescence imaging could present the targeted vessels above the liver with a higher SBR of 1.16 while the SBRs of the same vessel were 1.06 and 1.12 in fluorescence imaging beyond 1300 nm and 1500 nm. Thus, the results showed clearly that the extra collection of 1400-1500 nm was conductive to the precise deciphering and the NIR-IIx+NIR-IIb fluorescence detection would bring new ideas for off-peak fluorescence imaging and even the imaging-guided surgery in clinic.

**Fig. 7.**
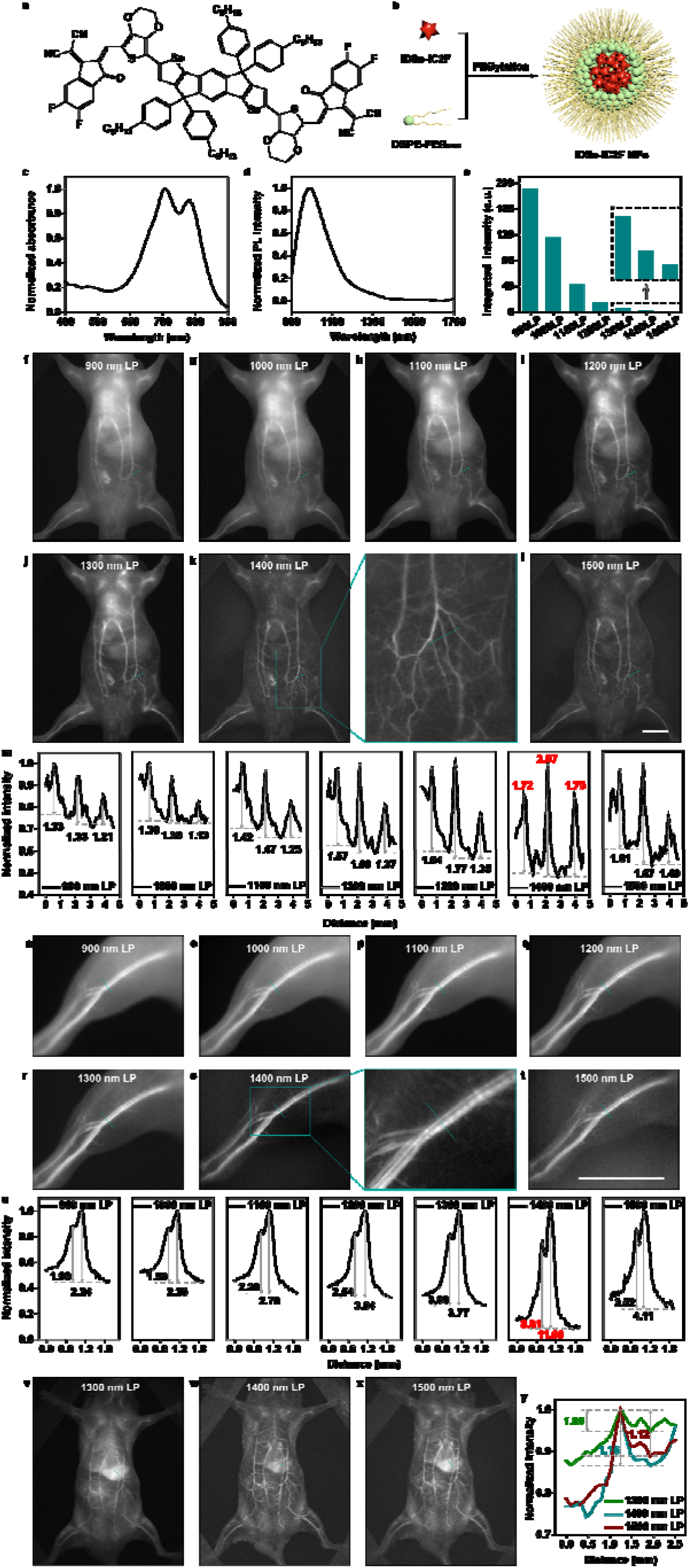
*In vivo* fluorescence imaging using the emission tail of the probes. **(a)** The molecular structure of IDSe-IC2F. **(b)** The schematic illustration of hydration. The normalized **(c)** absorbance and **(d)** emission spectra of IDSe-IC2F NPs. **(e)** The integrated emission intensity. The whole-body fluorescence imaging beyond **(f)** 900 nm, **(g)** 1000 nm, **(h)** 1100 nm, **(i)** 1200 nm, **(j)** 1300 nm, **(k)** 1400 nm and **(l)** 1500 nm. **(m)** Cross-sectional fluorescence intensity profiles along the indigo lines of the blood vessel in Fig. 7f-l. The numbers show the SBRs. The hind limb fluorescence imaging beyond **(n)** 900 nm, **(o)** 1000 nm, **(p)** 1100 nm, **(q)** 1200 nm, **(r)** 1300 nm, **(s)** 1400 nm and **(t)** 1500 nm. **(u)** Cross-sectional fluorescence intensity profiles along the indigo lines of the blood vessel in Fig. 7n-t. The numbers show the SBRs. The whole-body fluorescence imaging after the nanoparticles accumulated in the liver beyond **(v)** 1300 nm, **(w)** 1400 nm and **(x)** 1500 nm. **(y)** The fluorescence intensity profiles along the indigo lines of the blood vessel above the liver in Fig. 7v-x. The numbers show the SBRs. Scale bar, 10 mm.

## Discussion

Previous viewpoints have always emphasized that the decrescent photon scattering brought by the wavelength increase mainly contributed to the ameliorated performance of NIR-II fluorescence imaging but the light absorption peak by water has long been kept off. Admittedly, the excessive absorption of the excitation light might induce serious photo-thermal damage. However, the intensity of excited fluorescence is always far less than that of excitation light, thus the induced thermal effect could be ignored when emission region located near the absorption peak. On the other hand, it is generally accepted that the light absorption would also significantly decrease the useful signals. As for point detection (common in scanning microscopy), the excited pointed signals are collected into the detector (such as photomultiplier tube) and then the ballistic and diffused signals are both restructured on one pixel. On account of the effective signal collection, the absorption enhancement is usually undesirable in this situation. Notwithstanding, the signal diffusion is influential in the 2D area detection and our simulation results revealed that the moderate light absorption by the bio-tissue could attenuate the scattering photons with long optical path and thus restrain the image background in the 2D detection. Based on this, we defined the promising regions of 1400-1500 nm, 1700-1880 nm and 2080-2340 nm as NIR-IIx, NIR-IIc and NIR-III windows, and the NIR-III imaging was believed to provide the best imaging quality in the whole NIR region. We look forward to the novel fluorophores and the highly efficient detection technique in the future eagerly.

After verification of *in vivo* imaging, we believed that fluorescence imaging in the whole 900-1880 nm region showed desirable imaging quality resulting from the positive influence of incremental light absorption and decrescent photon scattering, compared with the imaging in the first near-infrared (NIR-I, 760-900 nm) window, the NIR-II region should be defined as 900-1880 nm, therefore. Besides, the promising performance of fluorescence imaging beyond 1300 nm, which was generally considered as NIR-IIa fluorescence imaging, further confirmed the dominance of the light absorption. NIR-IIx region, which was defined as 1400-1500 nm here, contained the intense absorption at ~1450 nm and were proved to own the optimum imaging potential, even exceeding the NIR-IIb region. When the emission is strong enough to resist the absorption loss, the collection within NIR-IIx region should be a recommendable option.

The off-peak NIR-II fluorescence imaging was an efficient compromise, on account of the limited peak emission wavelength of many current fluorescent probes. There have been some bright fluorophores developed for off-peak NIR-IIa or NIR-IIb fluorescence imaging, however, the NIR-IIx region was neglected. The long-pass usage with NIR-IIx+NIR-IIb detection (1400 nm LP) were proved to enhance image contrast. Adding the NIR-IIx collection, meanwhile, relaxed the requirements of the fluorophores, which benefited the off-peak fluorescence detection. The conclusion might guide the future off-peak fluorescence bioimaging and promote the development of the NIR-II fluorescent probes.

The poor sectioning strength of wide-field microscopy was attributed to the interference by the scattering and defocus signals. When shifting the imaging window around the NIR-IIx region, the image contrast was remarkably promoted. The performance of NIR-IIx fluorescence wide-field tomography was even comparable with that of the scanning microscopy, while the latter usually possessed more complex optical setup and lower temporal resolution. We believe an ingenious and simplified technique for sectioning imaging was proposed here and more imaging agents emitting around the NIR-IIx region in the future can be anticipated.

## Methods

### Simulation of near-infrared imaging

Monte Carlo method was utilized to simulate the propagation of photons in biological tissues. The total thickness of the tissue used in the simulation was 4 mm, with length and width of 100 mm and 100 mm, respectively. we assumed that the refractive index of the tissue was 1.37, the scattering anisotropy factor was 0.9, and the refractive index of air was 1. In the simulation, absorption coefficient of water was considered as the tissue absorption coefficient, and the reduced scattering coefficient was calculated using the following formula: μ_s’ =_ 1.5/λ (mm^−1^). The fluorescence signal source was a line with a length of 40 mm, a width of 1 mm and a depth of 1 mm in the tissue, which would emit a total of 10^6^ photons in random directions. The detection plane with 400×400 pixels is parallel to the tissue surface, which is 1 mm above the tissue surface. After a signal photon was successfully escaped from the tissue, it fell on a certain pixel on the detection surface. A simulated image was obtained by integrating the signals of all outgoing photons. Some point light sources with random emitting direction were added at random locations in the tissue as noise. There were 10^6^ noise photons in each simulation.

### Synthesis and PEGylation of the PbS/CdS quantum dots

For all details about synthesis and PEGylation of the CSQDs, please see the Supplementary Information.

### Animal preparation

All experimental procedures were approved by Animal Use and Care Committee at Zhejiang University. All the experimental animals involved, including BALB/c nude mice (~20 g) and Institute of Cancer Research (ICR) mice (~20 g), were provided from the SLAC laboratory Animal Co. Ltd. (Shanghai, China) and were fed with water and food with a normal 12 h light/dark cycle.

### Functional fluorescence imaging *in vivo*

All the experiments and operations on mice were conducted under anaesthesia. After intravenous injection of PbS/CdS CSQDs (1 mg mL^−1^, 200 μL) or IDSe-IC2F NPs (1 mg mL^−1^, 200 μL), the whole-body vessel imaging was performed under the expanded 793 nm CW laser excitation. As shown in Fig. S10, the NIR-II signals were collected via a fixed focus lens with near-infrared antireflection (TKL35, Tekwin, China) with the purification by different filters (The 900 nm, 1000 nm, 1100 nm, 1200 nm, 1300 nm, 1400 nm and 1500 nm long-pass filters were purchased from Thorlabs; The 1450 nm band-pass filter, 1550 nm long-pass filter, 1100 nm and 1300 nm short-pass filters were purchased from Edmund Optics; The 1550 nm short-pass filter was custom-built.) and eventually imaged on the electronic-cooling InGaAs camera (SD640, Tekwin, China). Next, the image distance was elongated to shrink the line field of view and thus the hind limb could be clearly presented. As for uterine imaging, the PbS/CdS CSQDs (1 mg mL^−1^, 200 μL) were infused into the uterine cavity via 26 G catheter. The mice with the uterus labeled were then placed in the macro imaging system and the bright NIR-II fluorescence from uterus could be further detected. To achieve NIR-II fluorescence cystography, the PbS/CdS CSQDs (1 mg mL^−1^, ~20 μL) were retrogradely injected into the bladder of the mouse with another 26 G catheter, then the bladder-labeled mice were moved to the macro imaging system.

### Wide-field microscopic fluorescence imaging *in vivo*

The skulls of the ICR mice were opened before imaging and the round thin coverslips were mounted on the brain. The treated mice were fixed after injected of the nanoparticles. As shown in Fig. S11, the 793 nm CW laser was introduced into the upright NIR-II fluorescence microscope (NIR II-MS, Sunny Optical Technology). The expanded 793 nm light was reflected by the dichroic mirror, passed through the objective and evenly excited the fluorophores covered in the bio-tissue. The fluorescence signals were collected by the objective, transmitted through the equipped filters and focused by the tube lens onto the imaging detector. Different magnification could be obtained by changing the objective.

## Acknowledgement

This work was supported by National Natural Science Foundation of China (61975172, 82001874, 61735016 and 21974104), Fundamental Research Funds for the Central Universities (2020-KYY-511108-0007) and Zhejiang Provincial Natural Science Foundation of China (LR17F050001). We acknowledge Dr. Tingting Luo (the Nanostructure Research Center, the Center for Materials Research and Analysis of Wuhan University of Technology) for the help in TEM analysis.

## Notes

### Competing Interest Statement

The authors have declared no competing interest.

