## supplementary information for "Perfecting and extending the near-infrared biological window"

#### Supplementary methods

**Materials and Methods Materials.**

$PbCl_2$, CdO, and sulfur powderwere purchased from Alfa. Oleylamine, oleic acid, 1-octadecene (ODE), tetrachloroethylene (TCE), poly(acrylic acid) (MW~1,800), N,N'-dicyclohexylcarbodiimide (DCC), N-(3-dimethylaminopropyl)-N′-ethylcarbodiimide hydrochloride (EDC), 2-(N-morpholino)ethanesulfonic acid (MES) were purchased from Sigma-Aldrich. Phosphate-buffered saline (PBS) was purchased from Hyclone. mPEG-amine (MW~5K) was purchased from Laysan Bio. 8-Arm PEG-amine (MW~40K) was purchased from Advanced BioChemicals. Other reagents were of analytical grade. Deionized ultra-filtered (DIUF) water was used in all needed experiments.

**Synthesis of PbS/CdS QDs.**

Sulfur precursor solution was prepared by mixing 0.08 g of sulfur powder, and 7.5 mL of oleylamine in two-neck flask at 120 ℃ under argon for 30 min. Lead precursor solution 0.834 g of $PbCl_2$ and 7.5 mL of oleylamine in a three-neck flask and degassing for 30 min under argon and heated to 45-150 °C (depending on the desired particle size). 2.25 mL of the sulfur precursor solution was injected into the Pb precursor solution under stirring. When the desired growth time is reached (typically 3-60 min), the reaction was quenched by adding 10 mL of cold hexane. The products were collected by centrifugation and re-suspended in 10 mL hexane/20 mL oleic acid. The mixture was agitated for 10 min to remove excess sulfur from the products. The QDs were precipitated via centrifugation. This precipitation procedure with oleic acid was repeated 3 times until the supernatant was colorless. After centrifugation of the suspension and decantation of the supernatant, the QDs were re-suspended in toluene/ODE.

The PbS/CdS CSQDs were synthesized via cation-exchange procedure. CdO (0.6 g, 4.6 mmol), oleic acid (4 mL), and ODE (15 mL) were heated to 200 °C, purged with Ar, and then cooled down to 100 °C. 5 mL of previously prepared PbS QDs suspended in Toluene/ODE was bubbled with Ar for 5 min, and then injected into the Cd precursor. The reaction flask was quenched with stopping stirring and heating directly after the growth reaction was conducted at 100 °C for 20-100 min. PbS/CdS CSQDs were precipitated with ethanol and then re-dispersed in hexane.

**Modification of CSQDs with oleyamine-branched polyacrylic acid (OPA).**

0.9 g of poly (acrylic acid) powder (average MW~1800) and 1.56 g DCC were transferred into a round-bottom flask. 10 mL of DMF was added to dissolve the mixture. About 1.2 mL of oleylamine were added dropwise into the reaction flask. The molar ratio of oleylamine to PAA is 30%. The solution was stirred overnight. 50 mL of 0.5 M HCl were added to the reaction solution. The precipitate was separated by centrifugation and re-dissolved in 3 mL methanol solution. And then 20 mL 1 M HCl was added to the solution. The precipitate was separated by centrifugation. This procedure was repeated at least 5 times. The precipitate was dissolved in 5 mL chloroform and washed by 10 mL 1 M HCl. The organic phase was collected and dried over by anhydrous $Na_2SO_4$. The chloroform was removed under vacuum, and the white solid was collected. The average Mw of OPA was ~ 3000 determined by gel permeation chromatography.

For surface modification, as-synthesized QDs (5.0 mg) were dissolved in 2.0 mL chloroform containing 15 mg of OPA. The mixture was stirred at room temperature for 30 min and the solvent was removed under vacuum by a rotary evaporator. The residue was then dissolved in 2 mL of 50 mM sodium carbonate solution under the sonication. The CSQDs were precipitated with ultracentrifuge at 750,00 rpm for 1 h. The purified product was dissolved in pH 8.5 MES buffer (0.01 M) and stored at 4 °C.

**PEGylation of PbS/CdS QDs.**

OPA-modified PbS/CdS QDs (5 mg) were dissolved in 20 mL pH 8.5 MES buffer (0.01 M). 15 mg of mPEG-amine (MW ~ 5K) and 5 mg of 8-Arm PEG-amine (MW ~ 40 K) with a molar ratio of 24: 1 were dissolved 1 mL MES, and gradually added to QDs solution with stirring. 10 mg of EDC was dissolved in 500 uL MES, and gradually added to QDs solution with stirring. The mixture was stirred at room temperature overnight. The PEGylated QDs was purified by 100 kDa filter, and washed five times with 1x PBS to remove excess reactants. The purified product was dissolved in 1x PBS and stored at 4 °C.

### Supplementary figures

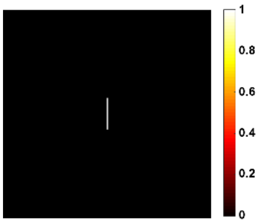


**Fig. S1 |** The line sample in simulation.

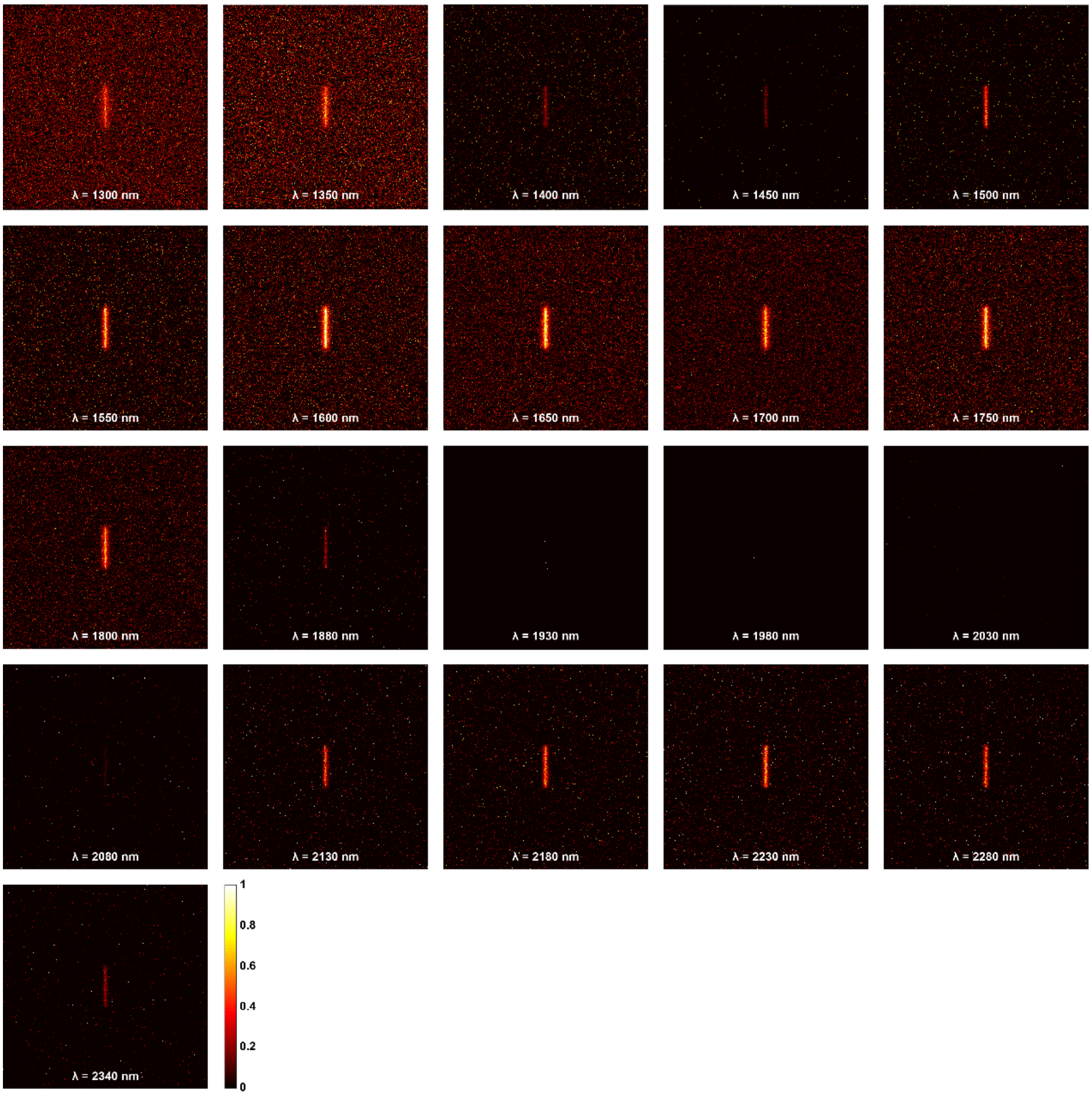


**Fig. S2 |** The simulated images at specific wavelengths.

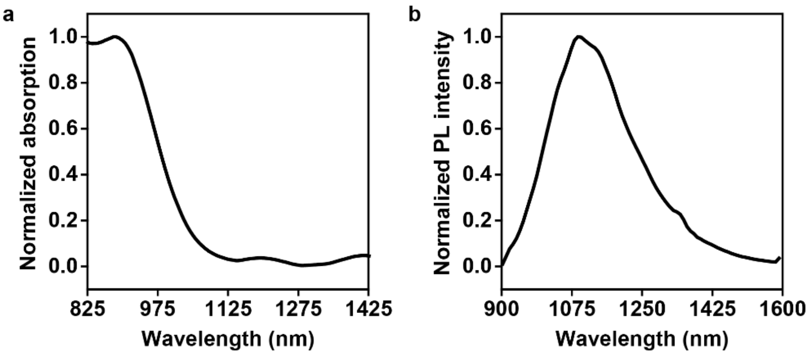


**Fig. S3 |** The normalized absorption (825-1425 nm) and PL intensity spectra of 1100-PbS/CdS QDs in tetrachloroethylene.

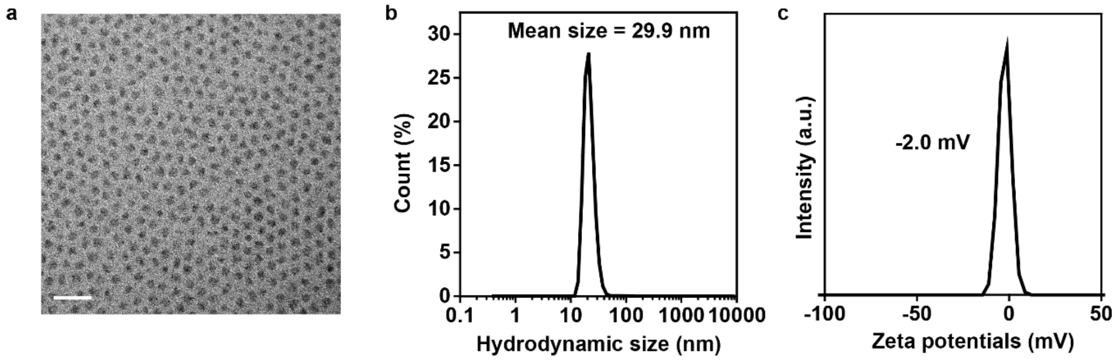


**Fig. S4 | (a)** The TEM image of as-prepared 1100-PbS/CdS QDs. The **(b)** hydrodynamic light scattering **(c)** and zeta potential results of the PEGylated 1100-PbS/CdS QDs.

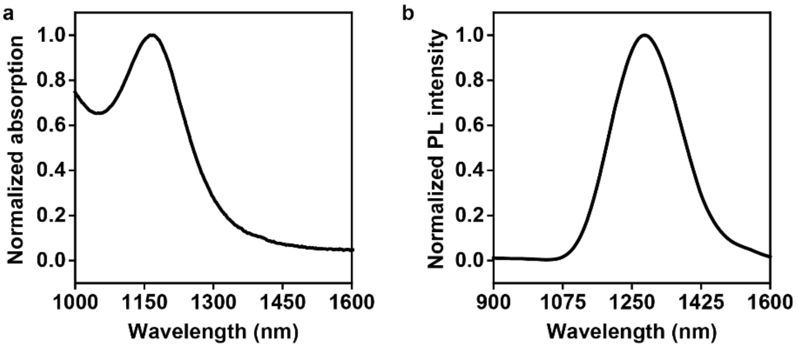


**Fig. S5 |** The normalized absorption (1000-1600 nm) and PL intensity spectra of 1300-PbS/CdS QDs in tetrachloroethylene.

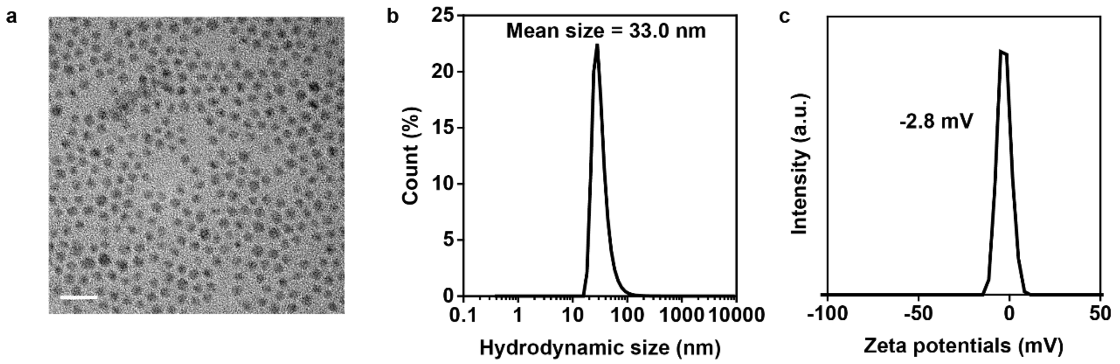


**Fig. S6 | (a)** The TEM image of as-prepared 1300-PbS/CdS QDs. The **(b)** hydrodynamic light scattering and **(c)** zeta potential results of the PEGylated 1300-PbS/CdS QDs.

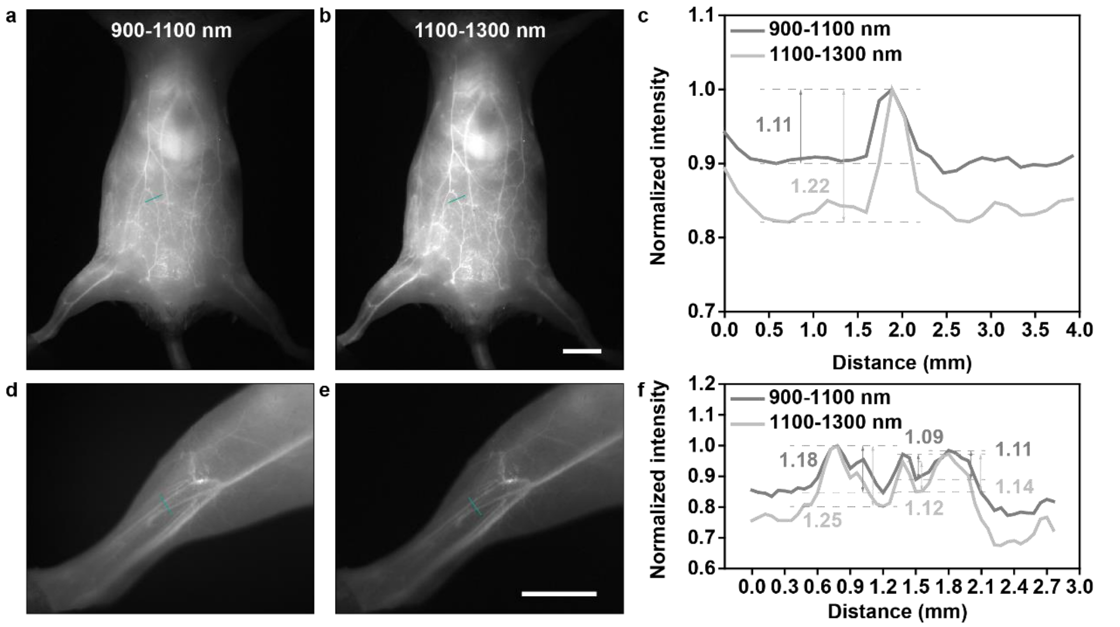


**Fig. S7 |** The whole-body imaging in **(a)** 900-1100 nm and **(b)** 1100-1300 nm. **(c)** Cross-sectional fluorescence intensity profiles along the indigo lines of the blood vessel in Fig. S7a & b. The numbers show the SBRs. The hind limb imaging in **(d)** 1100-1300 nm and **(e)** 1300-1500 nm. **(f)** Cross-sectional fluorescence intensity profiles along the indigo lines of the blood vessel in Fig. S7d & e. The numbers show the SBRs.

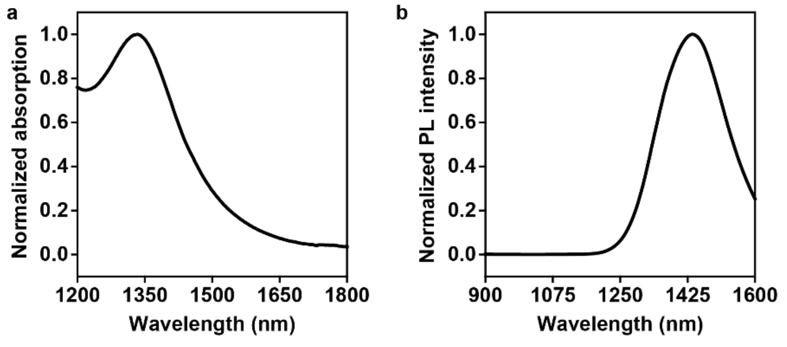

**Fig. S8 |** The normalized absorption (1200-1800 nm) and PL intensity spectra of 1450-PbS/CdS QDs in tetrachloroethylene.

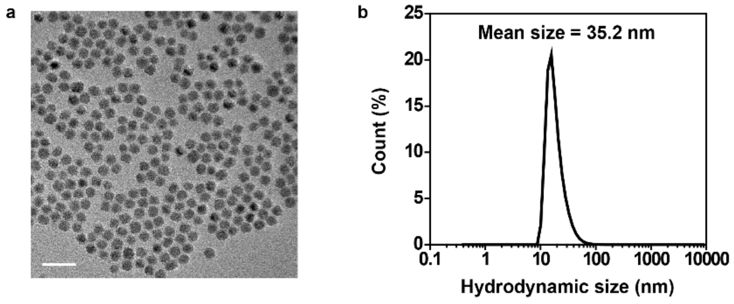


**Fig. S9 | (a)** The TEM image of as-prepared 1450-PbS/CdS QDs. The **(b)** hydrodynamic light scattering and **(c)** zeta potential results of the PEGylated 1450-PbS/CdS QDs.

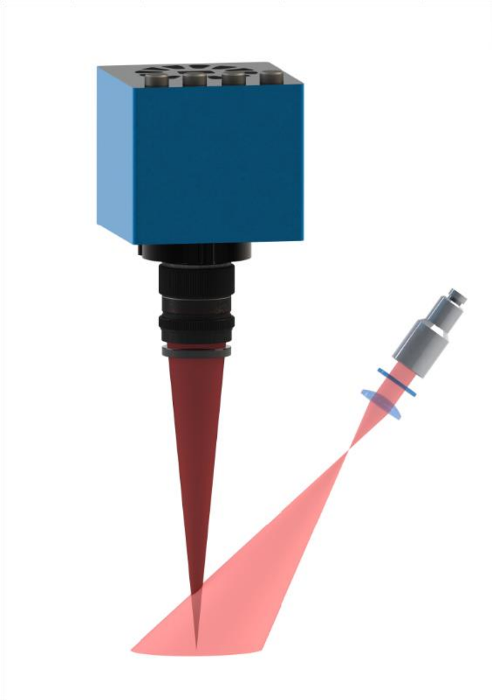

**Fig. S10 |** The macro imaging system.

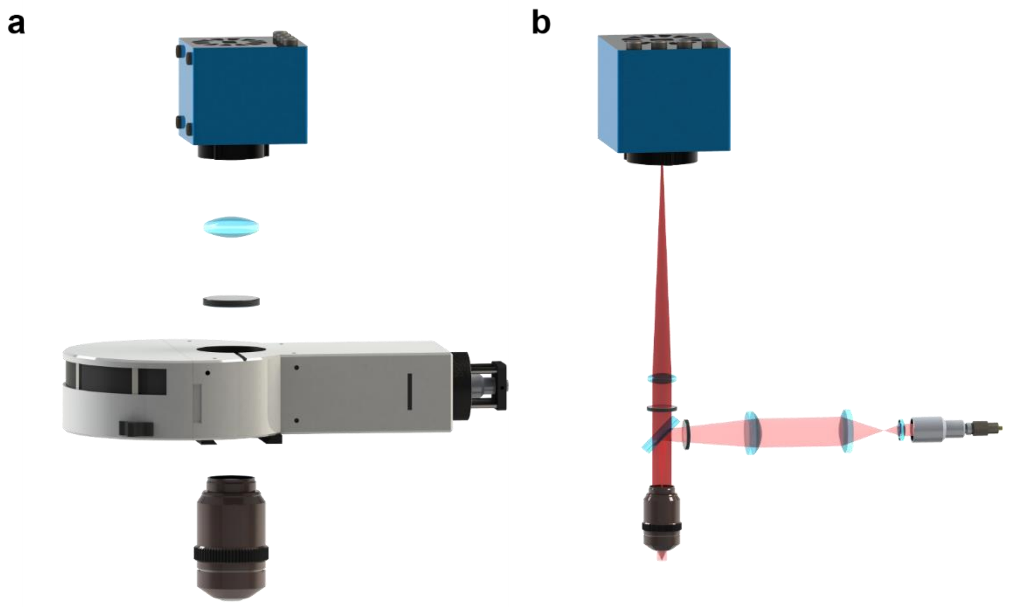


**Fig. S11 |** The **(a)** optical setup and **(b)** light-path diagram of the micro imaging system.
